# In Vitro Reconstitution of T Cell Receptor–Mediated Segregation of the CD45 Phosphatase

**DOI:** 10.1101/097600

**Authors:** Catherine B. Carbone, Nadja kerm, Ricardo A. Fernandes, Enfu Hui, Xiaolei Su, K. Christopher Garcia, Ronald D. Vale

## Abstract

T cell signaling initiates upon binding of peptide-major histocompatibility complex (pMHC) on an antigen-presenting cell (APC) to the T cell receptor (TCR) on a T cell. TCR phosphorylation in response to pMHC binding is accompanied by segregation of the transmembrane phosphatase CD45 away from TCR-pMHC complexes. The kinetic segregation hypothesis proposes that CD45 exclusion shifts the local kinase-phosphatase balance to favor TCR phosphorylation. Spatial partitioning may arise from the size difference between the large CD45 extracellular domain and the smaller TCR-pMHC complex, although parsing potential contributions of extracellular protein size, actin activity, and lipid domains is difficult in living cells. Here, we reconstitute segregation of CD45 from bound receptor-ligand pairs using purified proteins on model membranes. Using a model receptor-ligand pair (FRB-FKBP), we first test physical and computational predictions for protein organization at membrane interfaces. We then show that the TCR-pMHC interaction causes partial exclusion of CD45. Comparing two developmentally-regulated isoforms of CD45, the larger R_ABC_ variant is excluded more rapidly and efficiently (~50%) than the smaller R_0_ isoform (~20%), suggesting that CD45 isotypes could regulate signaling thresholds in different T cell subtypes. Similar to the sensitivity of T cell signaling, TCR-pMHC interactions with Kds of ≤15 μM were needed to exclude CD45. We further show that the co-receptor PD-1 with its ligand PD-L1, immunotherapy targets that inhibit T cell signaling, also exclude CD45. These results demonstrate that the binding energies of physiological receptor-ligand pairs on the T cell are sufficient to create spatial organization at membrane-membrane interfaces.

**SIGNIFICANCE STATEMENT:** The interface between a T cell and an antigen-presenting cell (APC) results in the formation of biochemically distinct plasma membrane domains that initiate signaling cascades. Here, using biochemical reconstitution and microscopy, we show that the binding energies of the TCRpMHC and PD-1-PD-L1 complexes are sufficient to create spatial organization at a model membrane-membrane interface. We show that spatial organization depends upon receptor-ligand binding affinity and the relative sizes of the extracellular domains. These biophysical parameters may be used to fine-tune signaling cascades in T cells.

## INTRODUCTION

Binding of the T cell receptor (TCR) to agonist peptide-Major Histocompatility Complex (pMHC) triggers a signaling cascade within a T cell leading to reorganization of the cytoskeleton and organelles, transcriptional changes, and cell proliferation. The first step in the cascade is TCR phosphorylation by the Src family tyrosine kinase Lck [Brownlie and Zamoyska, 2013]. One model, called “kinetic segregation” [Davis and van der Merwe, 2006], for how this initiating phosphorylation is triggered proposes that the close membrane contact created by TCR-pMHC binding results in exclusion of the transmembrane phosphatase CD45, and the shift of the kinase-phosphatase balance favors net phosphorylation of the TCR by Lck. The basis of this exclusion is thought to be steric, since the large CD45 extracellular domain (CD45 R_0_ isoform, 25 nm; CD45 R_ABC_ isoform, 40 nm, **Table S1**) [Woollett, et al., 1985; McCall, et al., 1992; Chang, et al. 2016] may not be able to penetrate the narrow inter-membrane spacing generated by TCR-pMHC complex (13 nm, **Table S1**) [Birnbaum, et al 2014; Choudhuri, et al 2005].

Imaging T cells activated *ex vivo* either by B cells [Leupin, et al. 2000] or by antigen presented on supported lipid bilayers [Johnson, et al., 2000; Varma, et al., 2006] has revealed that CD45 is indeed partitioned away from TCR upon pMHC binding. Cellular reconstitutions have demonstrated that the large extracellular domain of CD45 is required for this segregation [James and Vale, 2012; Cordoba, et al., 2013]. Additionally, size-dependent segregation of CD45 by orthogonal receptor-ligand pairs that create a similar narrow inter-membrane cleft, in the absence of TCR-pMHC binding, is sufficient for T cell triggering [James and Vale, 2012; Chang, et al., 2016].

Despite this strong cellular evidence for size-based partitioning, it has been debated whether the physical properties of CD45 and TCR-pMHC at the membrane-membrane interface alone are sufficient to explain the observed segregation behavior or whether other cellular factors (e.g. actin cytoskeletal or lipid ordering) are also required. Several groups have computationally modeled aspects of size-based organization at membrane interfaces, and two independent mathematical approaches have concluded that spontaneous pattern formation can occur in physiological parameter ranges [Lee et al., 2003; Weikl and Lipowsky, 2004]. These models predict the contributions of protein (size, concentration, elasticity, affinity and kinetics), membrane (stiffness, tension, repulsion), and environmental factors (thermal fluctuations, cytoskeleton, time) in regulating partitioning. Although these models focus primarily on a system with two binding pairs (TCR-pMHC and ICAM-1-LFA-1), some of the predictions can be extrapolated to a system with both ligand-bound and unbound species.

Successful efforts to reconstitute molecular segregation at membrane-membrane interfaces have been made with dimerizing GFP molecules [Schmid, et al., 2016] and hybridizing strands of DNA [Chung, et al., 2013]. These studies show that laterally mobile molecules at membrane-membrane interfaces organize by height and locally deform the membrane to accommodate different molecular sizes. However, results from high affinity, artificial receptor-ligand pairs cannot be simply extrapolated to predict results for physiologically relevant molecules at the T cell-APC interface. Here, we have recapitulated TCR-pMHC-mediated partitioning of CD45 on model membranes.

## RESULTS

### A chemically-inducible receptor-ligand system for producing CD45 exclusion at a membrane-membrane interface

To mimic a T cell, we used a giant unilamellar vesicle (GUV) containing a nickel-chelating lipid to which a purified His_10_-tagged, fluorescently-labeled receptor and CD45 could be added (**Fig. 1A**). To mimic the APC, we used a supported lipid bilayer (SLB) containing nickel-chelating lipids to which a His_10_-tagged protein ligand also could be bound. As an initial test of this system, we used an artificial receptor (FKBP) and ligand (FRB) that could be induced to form a tight binding interaction (100 fM) upon addition of rapamycin [Banaszynski, et al., 2006]. In order to maintain the GUV and SLB in proximity prior to rapamycin addition, the two membranes were passively tethered to one another using two 100-mer single-stranded DNA molecules with a 20 bp region of complementarity [Taylor et al., 2017, Chi et al., 2013] (**Table S1**). The elongated extracellular domain of the CD45 R_0_ isoform (25 nm) [Woollett, et al., 1985; McCall, et al., 1992; Chang, et al. 2016] or the smaller SNAP protein (5 nm, **Table S1**) [Gautier, et al. 2008] were used as test proteins for partitioning.

**Fig. 1.**
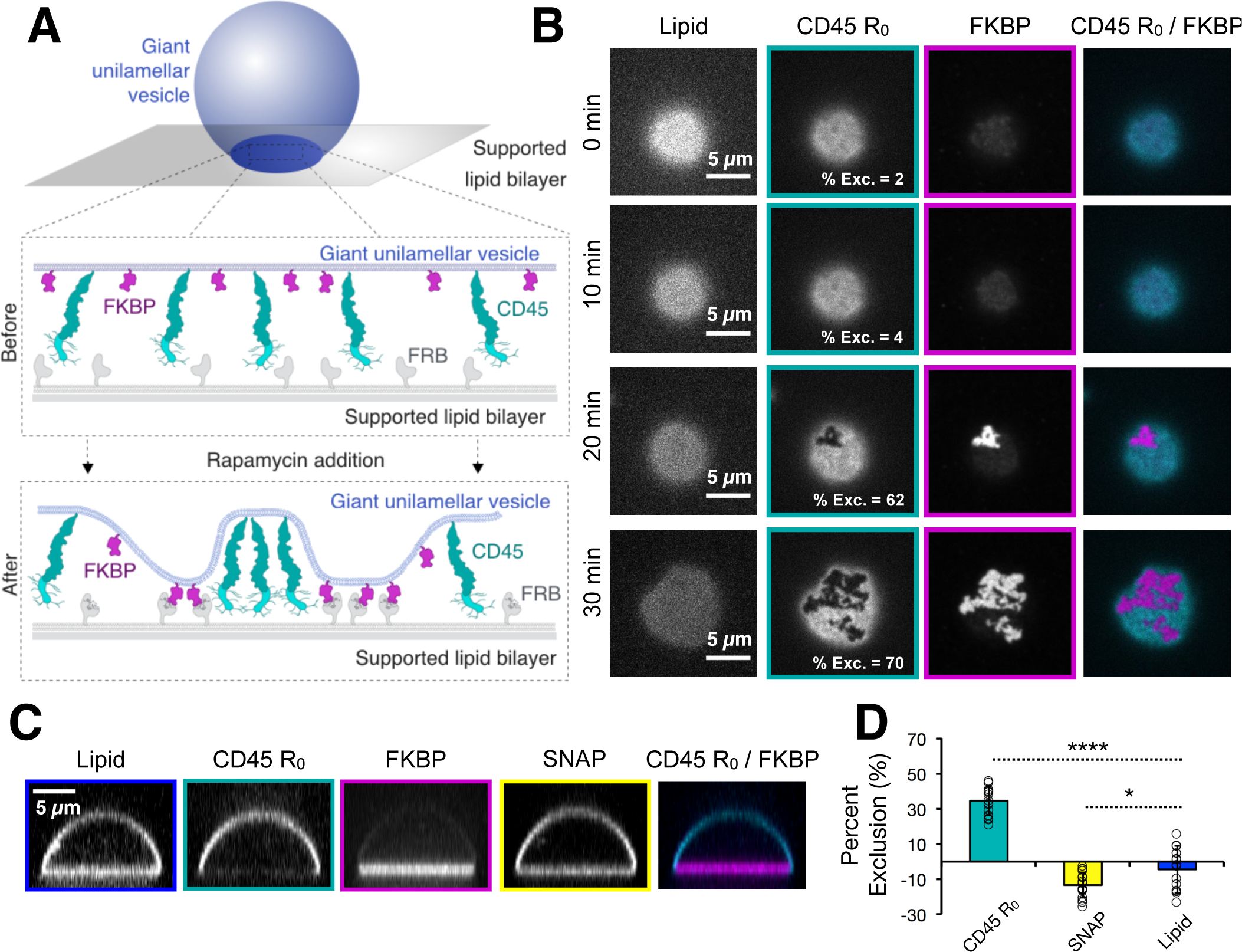
Receptor-ligand binding induces CD45 segregation at membrane interfaces. (**A**) Schematic of rapamycin-induced receptor (FKBP)-ligand (FRB) binding and CD45 R_0_ segregation between a giant unilamellar vesicle (GUV) and a supported lipid bilayer (SLB) (**B**) Total internal reflection fluorescence (TIRF) microscopy of a GUV-SLB interface at indicated times after rapamycin addition, showing concentration of FKBP into microdomains that exclude CD45 R_0_. Percent exclusion of CD45 R_0_ is indicated for each image shown. (**C**) Spinning disk z-sections of GUVs after membrane-apposed interfaces have reached equilibrium, showing localization of FKBP to the membrane interface, localization of CD45 R_0_ away from the interface, and uniform distribution of SNAP. (**D**) Quantification of experiment shown in **C**; mean ± standard deviation (n=17 GUVs pooled from two experiments).

Upon rapamycin addition, FKBP and FRB concentrated first in small micron-scale clusters at the GUV-SLB interface, which then grew in size over the interface; simultaneously, fluorescently-labeled CD45 R_0_ partitioned away from regions of the GUV that became enriched in receptor-ligand (**Fig. 1B and Movie S1**). In contrast to CD45, which was strongly depleted by FRB-FKBP, the SNAP protein (5 nm) [Schmitt, et al 2010] or a lipid dye (Atto390-DOPE) remained evenly distributed throughout the interface after rapamycin addition (**Fig. 1C-D**). The size of FKBP-FRB clusters could be varied by changing the receptor concentration on the GUV membrane; however, the degree of CD45 R_0_ exclusion from clusters was similar over the range tested (**Fig. 2A-C**). Across all concentrations of FKBP, at receptor-ligand enriched zones, CD45 R_0_ was depleted by 72 ± 7% (n=22 GUVs pooled from two experiments). Once formed, the receptor -enriched and -depleted zones stably retained their shapes for tens of minutes. However, using single molecule TIRF imaging, we observed that single molecules of CD45 R_0_ can diffuse across FKBP-FRB -enriched and -depleted zones (**Fig. 2D-E, Movie S2**). This result reveals that individual molecules can exchange across these micron-scale boundaries.

**Fig. 2.**
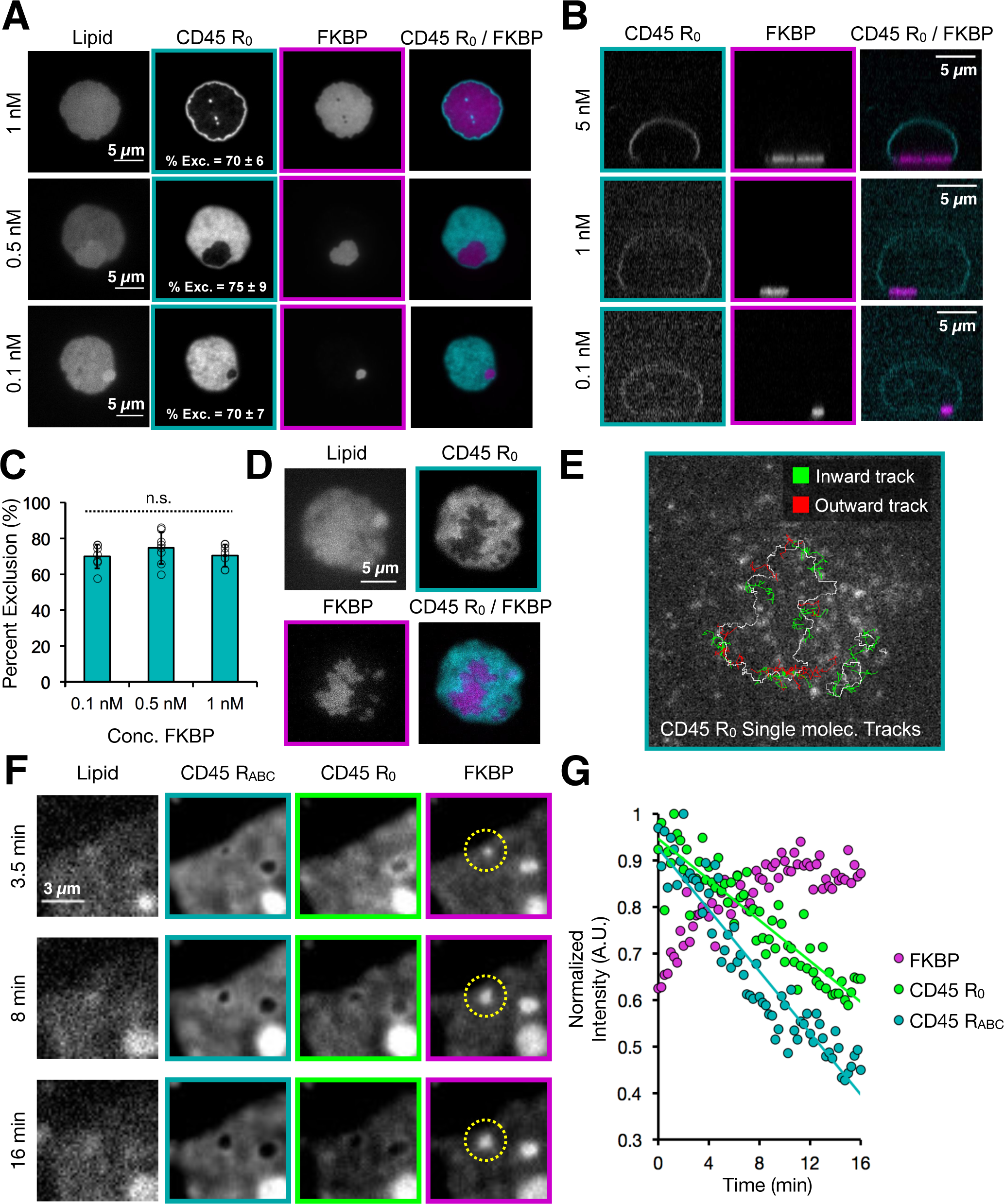
Characterization of partitioned GUV-SLB membrane-membrane interfaces. (**A**) Titration of FKBP concentration (indicated at left of images) with constant CD45 R_0_ concentration imaged by TIRF microscopy. Percent exclusion of CD45 R_0_ is indicated as mean ± standard deviation with n=7-8 GUVs per condition pooled from three experiments. (**B**) Spinning disk z-sections of GUVs shown in **A**. (**C**) Graphical representation of data shown in **A**. (**D**) Total internal reflection fluorescence (TIRF) microscopy of a GUV-SLB interface showing overall localization of CD45 R_0_ and FKBP. (**E**) Single molecule imaging of CD45 R_0_ for GUV shown in **D**, border of FKBP enriched zone indicated by white line. Only tracks crossing the exclusion boundary are shown. CD45 R_0_ single molecule tracks originating outside FKBP enriched zone are shown as green lines and tracks originating inside the FKBP enriched zone are shown as red lines. (**F**) Total internal reflection fluorescence (TIRF) microscopy of a GUV-SLB interface at 30-sec time points after rapamycin addition showing concentration of FKBP into micro domains that exclude CD45 R_0_ and CD45 R_ABC_. Rate of CD45 R_ABC_ exclusion is 2.8 ± 0.9 times faster than rate of CD45 R_0_ exclusion, n=7 GUVs from two experiments. (**G**) Quantification of exclusion for representative GUV shown in **F**.

In addition to testing the CD45 R_0_ isoform for segregation, we also compared the extracellular domain of the CD45 R_ABC_ isoform, which is preferentially expressed early in T cell development [Hermiston, et al., 2003], and is about 15 nm larger in size than the shorter and later expressed R_0_ isoform (**Table S1**) [Woollett, et al., 1985; McCall et al., 1992]. With both isoforms present on the same GUV, the larger CD45 R_ABC_ isoform segregated from newly forming FKBP clusters three-fold faster than the R_0_ isoform (2.8 ± 0.9-fold, n=7 GUVs pooled from two experiments, **Fig. 2F-G, Movie S3**). However, the final extent of exclusion between the two CD45 isoforms was similar with this high affinity FRB-FKBP system (**Fig. S2**).

The kinetic segregation model predicts that CD45 is excluded from receptor-ligand complexes based upon a difference in the spacing between the GUV and SLB in the receptor- versus CD45-enriched regions [Davis and van der Merwe, 2006]. To investigate the topology of the GUV membrane across the interface with nanometer accuracy in the vertical axis, we used scanning angle interference microscopy (SAIM), a technique that calculates the distance of fluorophores from a silicon oxide wafer by collecting sequential images at multiple illumination angles (**Fig. 3A**) [Carbone, et al., 2016]. The SAIM reconstructions revealed membrane deformations at regions of CD45 localization (**Fig. 3B-D**). The calculated difference in membrane spacing between the FRB-FKBP- and CD45 R_0_- enriched regions was 18 ± 11 nm (n=4-6 regions from each of 4 GUVs from two experiments, pooled), suggesting a size of ~24 nm for the CD45 R_0_ extracellular domain, assuming that FRB-FKBP creates an intermembrane space of 6 nm (**Table S1**)[Liang, et al., 1999]. This value is similar to the ~22 nm axial dimension for the CD45 R_0_ extracellular domain determined by electron microscopy (Chang et al., 2016). Conversely, for GUV-SLB interfaces with FRB-FKBP and SNAP, SAIM reconstructions revealed no changes in membrane spacing across the GUV-SLB interface (**Fig. 3E-G**).

**Fig. 3.**
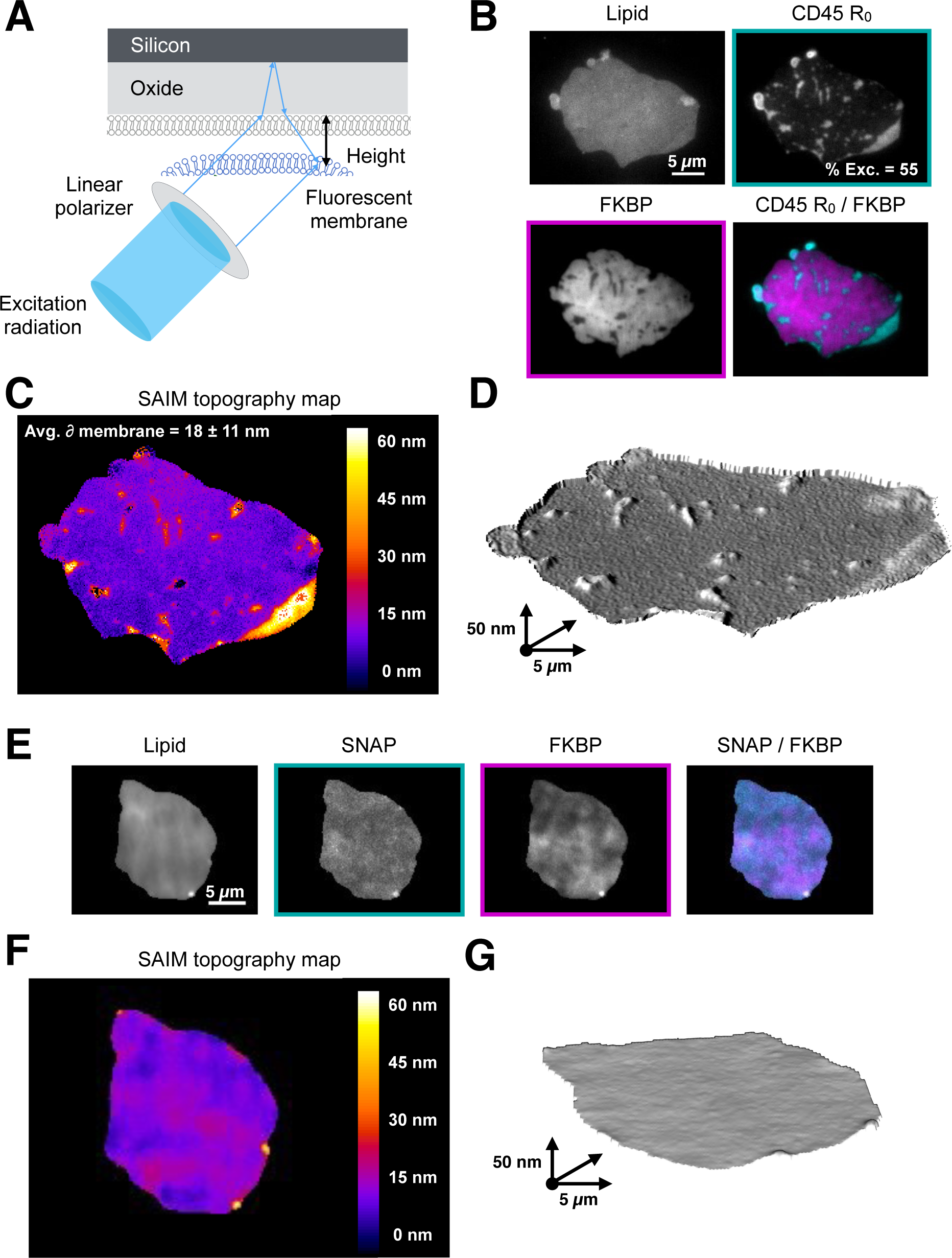
Membrane topology is influenced by local protein composition. (**A**) Schematic of scanning angle interference microscopy showing reflection and interference of excitation light that produces structured illumination patterns used to deduce fluorophore height; adapted from Carbone, et al., 2016. (**B**) Epifluorescence microscopy showing localization of lipid, CD45 R_0_ and FKBP on GUV analyzed by SAIM imaging. Percent exclusion of CD45 R_0_ indicated for image shown. (**C**) SAIM reconstruction of GUV membrane derived from lipid fluorescence showing an increase in membrane height at CD45 R_0_ clusters. Average membrane height change depicted as mean ± standard deviation, n=4-6 clusters from each of 4 GUVs imaged during two separate experiments. (**D**) 3D model of data shown in **c**. Z-scale is exaggerated to clearly depict membrane deformations. (**E**) Epifluorescence microscopy showing localization of lipid, SNAP, and FKBP on GUV analyzed by SAIM imaging. (**F**) SAIM reconstruction of GUV membrane derived from lipid fluorescence (**G**) 3D model of data shown in **F**. Z-scale is exaggerated to clearly depict membrane deformations.

### TCR-pMHC –mediated CD45 exclusion

Next, we sought to establish a GUV-SLB interface using the native T cell receptor-ligand pair, TCR-pMHC (**Fig. 4A**). For the TCR, we co-expressed the extracellular domains of the 2B4 α and β chains extended with leucine zippers to stabilize their dimerization [Birnbaum et al., 2014]; both chains were tagged with His_10_ for conjugation to the GUV membrane and the β chain contained a ybbR peptide for fluorescent labeling. For the ligand, we used the IE^k^ MHC, His_10_-tagged loaded with a high affinity (2.5 μM Kd) peptide. Similar to the results previously described for FRB-FKBP, we observed the formation of micron-sized TCR clusters that excluded CD45 R_0_ (22 ±14% exclusion, n=17 GUVs pooled from 2 experiments, **Fig. 4B**) but not the control SNAP domain (**Fig. S2A**).

**Fig. 4.**
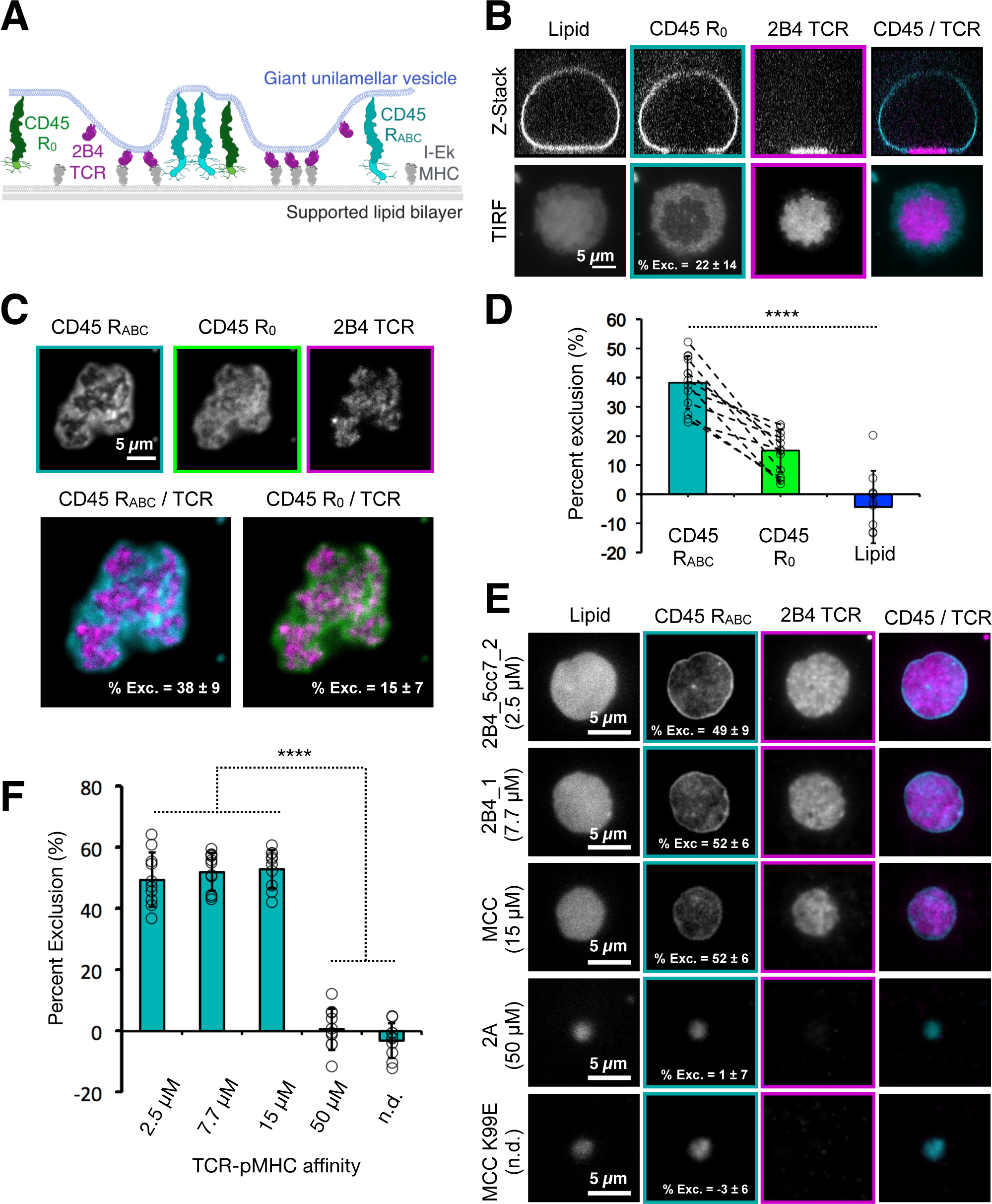
TCR-pMHC binding induces CD45 segregation at GUV-SLB interfaces. (**A**) Schematic of 2B4 TCR-IE^k^ MHC binding between a GUV and a SLB, and segregating away from two CD45 isoforms (R_0_ and R_ABC_). (**B**) Top, spinning disk z-sections of GUVs after membrane-apposed interfaces have reached equilibrium, showing localization of 2B4 TCR to membrane interface and exclusion CD45 R_0_ away from the interface. Bottom, TIRF images of GUV-SLB interface for GUV shown in panel above. Percent exclusion of CD45 R_0_ indicated for image shown. (**C**) Top, segregation of CD45 R_0_ and CD45 R_ABC_ on the same GUV membrane away from 2B4 TCR, shown by TIRF microscopy of membrane interface. Percent exclusion of CD45 isoforms indicated as mean ± standard deviation, with n=13 GUVs from two experiments. (**D**) Graphical representation of data shown in **C**. (**E**) Dependence of CD45 R_ABC_ exclusion as a function of TCR-pMHC affinity using peptides with different Kds, indicated at left of images. Imaged by TIRF microscopy of membrane interfaces. Percent exclusion of CD45 _RABC_ indicated as mean ± standard deviation, n=10 GUVs per condition from two experiments. (**F**) Graphical representation of data shown in **E**.

We also combined both CD45 R_ABC_ and CD45 R_0_ isoforms on the same GUV and compared their segregation with the TCR-pMHC system. Upon GUV contact with the SLB, the 2B4 TCR bound the IE^k^ MHC, and concentrated at the interface where it formed micron-scale clusters that excluded both isoforms of CD45 (**Fig. 4C**). However, unlike the high affinity FKBP-FRB system, the degree of TCR-pMHC mediated exclusion of the smaller CD45 R_0_ isoform (15 ± 7% exclusion) was lower than the larger CD45 R_ABC_ isoform (38 ± 9% exclusion) at steady state (45 min, n=13 GUVs pooled from two experiments, **Fig. 4D**).

*In vivo*, TCR encounters MHCs loaded with a myriad of different peptides; although not absolute, TCR-pMHC affinities of <50 μM are usually required to trigger a signaling response [Gascoigne, et al., 2001]. To examine the effect of TCR-pMHC affinity on CD45 R_ABC_ exclusion, we loaded IE^k^ MHC with a series of well-characterized peptides with resultant Kds of 2.5 μM, 7.7 μM, 15 μM, 50 μM and null for the 2B4 TCR [Birnbaum, et al., 2014]. At steady state, we observed that pMHCs with affinities to the TCR of 15 μM and lower excluded CD45 R_ABC_ to similar extents (51 ± 7% exclusion, n=30 GUVs pooled from two experiments, **Fig. 4E-F**). However, the pMHC with a Kd of 50 μM and IE^k^ loaded with null peptides did not concentrate TCR at the GUV-SLB interface and did not change the distribution of CD45 R_ABC_ (-1 ± 6% exclusion, n=20 GUVs pooled from 2 experiments, **Fig. 4E-F**). Thus, in agreement with computational predictions [Weikl and Lipowsky, 2004], CD45 R_ABC_ exclusion was observed over the same range of affinities that are associated with peptide agonists.

### Exclusion of CD45 by PD-1 –PD-L1

T cell signaling involves many receptor-ligand pairs interacting across the two membranes in addition to the TCR-pMHC [Chen and Flies, 2013]. The co-receptor PD-1 and its ligand PD-L1 create a signaling system that opposes T cell activation by inhibiting CD28 signaling [Hui et al, 2017; Kamphorst, et al, 2017]. PD-1 ligation also results in microcluster formation on T cells [Yokosuka st al, 2012]. Like the TCR, PD-1 signaling is initiated through receptor tail phosphorylation by Lck [Sheppard et al, 2004], and this phosphorylation event may be opposed by the abundant CD45 phosphatase (**Fig. S2A-B**). Therefore we tested the ability of interaction of PD-1 with PD-L1, which forms a complex of similar dimension (9 nm) to TCR-pMHC (**Table S1**) [Lin, et al., 2008]) to partition CD45 in our in vitro liposome system (**Fig. 5A**). As expected from these physical dimensions, PD-1-PD-L1 interaction at the membrane-membrane interfaces formed micron-sized clusters that excluded CD45 R_ABC_ (**Fig. 5B**). The CD45 R_ABC_ The degree of ABC exclusion (60 ± 14% exclusion, n=14 GUVs from two experiments **Fig. 5B**) was greater than that observed for TCR-pMHC (2.5 μM peptide), which may be explained by the higher affinity of the PD1-PD-L1 interaction (0.77 μM, [Butte, et al., 2013].

**Fig. 5.**
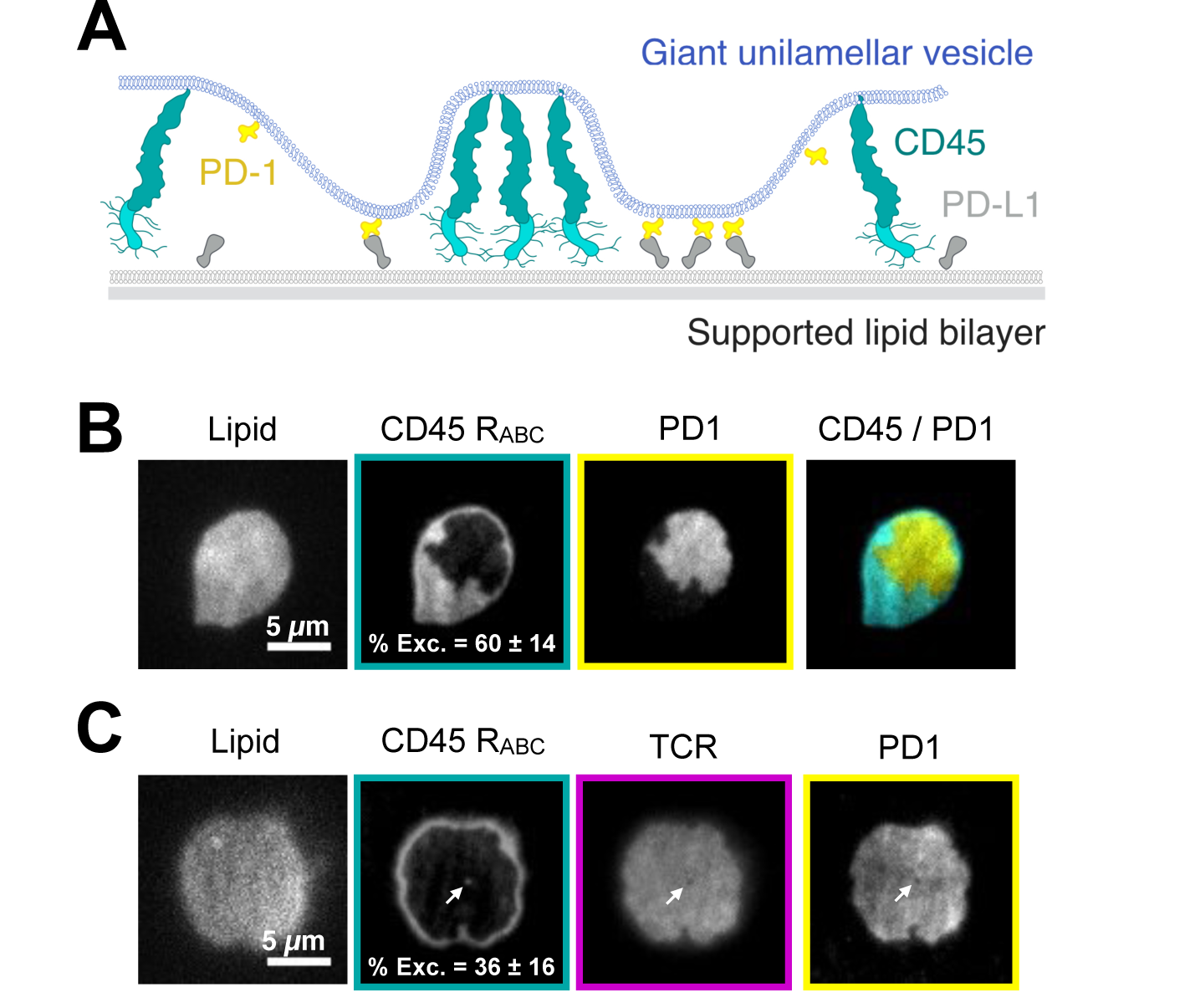
The inhibitory co-receptor PD-1 excludes CD45 and colocaizes with TCR. (**A**) Schematic of PD-1-PD-L1 binding between a GUV and a SLB, with segregation away from CD45 R_ABC_. (**B**) TIRF microscopy showing concentration of PD-1 into microdomains that exclude CD45 RABC. Percent exclusion of CD45 R_ABC_ indicated as mean ± standard deviation, n=14 GUVs from two experiments. (**C**) TIRF microscopy showing concentration of TCR and PD-1 into a domain that excludes CD45 RABC. Percent exclusion of CD45 R_ABC_ indicated as mean ± standard deviation, n=14 GUVs from two experiments. White arrow highlights small CD45 R_ABC_ enriched zone that is depleted for TCR and PD-1.

We also combined CD45 R_ABC_ with both TCR-pMHC with PD-1-PD-L1. In this dual receptor-ligand system, the two receptor-ligand complexes co-localized and CD45 R_ABC_ was partitioned away from the combined ligated TCR-PD-1 footprint (**Fig. 5C**). The size (**Table S1**) and affinity difference between TCR-pMHC and PD-1-PD-L1 may be small enough to not cause partitioning of these receptor-ligands under the conditions tested in our in vitro assay.

## DISCUSSION

In this study, we have established an *in vitro* membrane system that recapitulates receptor-ligand mediated CD45 exclusion. We have found that the binding energy of physiological receptor-ligand interactions is sufficient for CD45 partitioning at a model membrane-membrane interface. We also show that subtle differences in sizes and affinities of the proteins at the interface can give rise to significant changes in spatial organization and discuss the implications of these findings in more detail below.

Spatial organization of TCR and CD45 at the immune cell contacts has been proposed to arise by a nucleation-spreading mechanism [Weikl and Lipowsky, 2004]. By imaging an inducible synthetic receptor-ligand binding interaction in real time, we also conclude that pattern formation arises by the nucleation of small clusters that further spread across the membrane interface over time. These patterns induce changes in membrane topology that reflect the local protein composition and are stable on the order of hours. However, we show that individual molecules can freely exchange between domains. This result is consistent with previous computational simulations, although these models predict patterns will relax to a circular geometry to minimize the length of the domain boundaries [Burroughs and Wulfing, 2002; Weikl and Lipowsky, 2004, Krobath, et al., 2011]. In our system, as observed for other physical models of partitioning using DNA-DNA hybridization [Chung, et al., 2013] and dimerizing GFP [Schmid, et al., 2016], patterns have more complex domain structures. The lack of circular geometry in the experimental systems could be due to small inhomogeneities in the supported lipid bilayer compared to perfectly diffusive computational models. Despite this difference, many physical and computational model systems have converged on nucleation and spreading as a general mechanism by which spatial organization arises at membrane-membrane interfaces.

The mechanism by which receptor-ligand binding induces spatial organization is a subject of active investigation. Our results showing differential exclusion of CD45 R_0_ and CD45 R_ABC_ indicate that size-based steric exclusion and membrane deformation are important for exclusion. In addition, protein crowding of receptor-ligand complexes also could provide a driving force for partitioning. Indeed, previous work has shown that patterns formed at analogous membrane-membrane interfaces using dimerizing GFP as the receptor-ligand pair and a small test protein (monomeric Cherry) are due to crowding effects [Schmid, et al., 2016]. In our system, however, we observe that the small SNAP protein is distributed throughout receptor-ligand enriched and depleted zones. These systems employ different proteins at the interface, and it will be interesting to investigate whether specific protein properties (e.g. size, propensity for oligomerization, elasticity, flexibility, packing density of receptor-ligand in partitioned zones, etc) account for these differences in the role of protein crowding in exclusion.

Our work also suggests an important contribution of receptor-ligand affinity in protein exclusion. We observed 70% depletion of CD45 R_0_ from FRB-FKBP (100 fM Kd) -enriched zones. The TCR-pMHC interactions, on the other hand, are much lower in affinity, with most agonists generally displaying Kds of 1-100 μM [Gascoigne, et al., 2001]. Strikingly, when we tested CD45 exclusion using TCR-pMHC, we found that exclusion was only 27% for the R_0_ isoform and 49% for the R_ABC_ isoform when tested individually. The PD-1-PD-L1 interaction is higher affinity (0.7 μM) and produces a somewhat higher exclusion (60%) of CD45 R_ABC._ While the CD45 R_0_ isoform exclusion by TCR-pMHC is modest, it nevertheless could be significant for eliciting a signaling response. *In vitro* analysis of the kinase-phosphatase network controlling TCR activation has shown that at physiological protein densities, small perturbations of CD45 can drive large changes in TCR phosphorylation [Hui et al, 2014]. In combination with our results, this suggests that the cellular CD45 concentration may position the TCR precisely at the boundary of a switch-like response in phosphorylation.

Our experimental results also are in reasonable agreement with computational predictions for a lower boundary of receptor-ligand affinity needed for protein exclusion. Computational models by Weikl et al. [Weikl, et al., 2003] predict that, at the ratio of 1 TCR molecule to 8 CD45 molecules used in these experiments, a binding energy of >4 k_B_T (corresponding to a Kd of ~20 μM) is required for partitioning. In our system, we find that a pMHC ligand with 15 μM Kd causes CD45 exclusion whereas a ligand with a Kd of 50 μM does not. It also has been predicted that increasing the affinity of a receptor-ligand interaction should increase the area fraction of the interface occupied by the receptor-ligand enriched zone by increasing the number of bound complexes at the same protein densities [Weikl and Lipowsky, 2004; Chung et al 2013]. However, in our experiments, TCR-pMHC mediated CD45 partitioning occurs as an all-or-nothing process.

Our results also demonstrate that the large extracellular domains of CD45 R_ABC_ and CD45 R_0_ are differentially sensitive to the partitioning forces produced by ligand-receptor binding interactions at a membrane-membrane interface. This finding is consistent with results showing that T cells expressing larger CD45 isoforms signal more efficiently [Chui, et al., 1994], although others have contested this conclusion [Czyzyk, et al., 2000]. Although the signaling consequences of differential CD45 segregation on immune activation remain to be clarified, our results establish a biophysical difference between two highly conserved CD45 isoforms [Okumura, et al., 1996] with regard to their degree of spatial segregation in response to TCRpMHC interactions. Given that the smaller CD45 isoforms are preferentially expressed in later steps of T cell selection [Hermiston et al., 2003], our results suggest that T cell signaling may be attenuated by changes CD45 isoform expression as a mechanism of peripheral tolerance.

We also explore increasing complexity at a membrane interface by introducing two receptor-ligand pairs: TCR-pMHC and PD-1-PD-L1. Interestingly, we find that these two receptor-ligands complexes co-localize with one another and both together exclude CD45. *In vivo,* partial segregation of these two receptor-ligands also has been observed in CD8+ T cells (Hui et al., 2017), and a higher degree of co-localization between these receptors was reported in CD4+ T cells (Yokosuka et al., 2012). Given that the size difference between the TCR-pMHC and PD-1-PD-L1 lies at the biophysical threshold for partitioning [Schmid et al., 2016], these results suggest that cellular localization of PD-1 with respect to TCR may be regulated by other factors (e.g. other co-receptors or adaptor proteins) and perhaps even in cell type -specific manner. In addition, it will be interesting to investigate how actin polymer dynamics and lipid-mediated organization [Koster and Mayor, 2016] may enhance or disrupt protein patterning across two membranes.

## ACKNOWLEDGMENTS

We would like to thank N. Stuurman for help with microscopy and image analysis and M. Taylor for guidance with protein purification and DNA tethering. We thank A. Williamson, N. Stuurman, and M. Morrissey for comments on the manuscript. The authors acknowledge funding from the Howard Hughes Medical Institute and National Institutes of Health (R01EB007187, R.D.V.).

## COMPETING INTERESTS

The authors do not declare any competing interests.

## MATERIALS AND METHODS

### Materials

Synthetic 1,2-dioleoyl-sn-glycero-3-phosphocholine (POPC; Avanti, 850457), 1,2-dioleoyl-sn-glycero-3-[(N-(5-amino-1-carboxypentyl)iminodiacetic acid)succinyl] (nickel salt, DGS-NTA-Ni; Avanti, 790404) and 1,2-dioleoyl-sn-glycero-3-phosphoethanolamine-N [methoxy(polyethylene glycol)-5000] (ammonium salt, PEG5000-PE; Avanti, 880220) were acquired from Avanti Polar Lipids, Alabama, USA. 1,2-dioleoyl-sn-glycero-3-phosphoethanolamine-Atto390 (DOPE-390; AttoTec, AD390-161) was acquired from Atto-Tec, Germany.

### Recombinant protein expression, purification, and labeling

N-terminally His_10_- and SNAP-tagged FRB and FKBP were subcloned into a pET28a vector and were bacterially expressed in BL21(DE3) strain of *Escherichia coli*. The cells were lysed in an Avestin Emulsiflex system. C-terminally His_10_- and SNAP-tagged extracellular domains of human CD45 R_0_, human CD45 R_ABC_, and human PD-L1 were subcloned into a pFastBac vector and were expressed in SF9 cells. All proteins were purified by using a HisTrap excel column (GE Healthcare Life Sciences) following the product recommendations. Recombinant C-terminal His_10_-tagged mouse PD-1 extracellular domain was purchased from Sino Biological.

2B4 TCR V_m_C_h_ chimeras containing an engineered C domain disulfide were cloned into the pAcGP67a insect expression vector (BD Biosciences, 554756) encoding either a C-terminal acidic GCN4-zipper-Biotin acceptor peptide (BAP)-6xHis tag (for α chain) or a C-terminal basic GCN4 zipper-6xHis tag (for β chain) [Wilson, et al., 1999]. Each chain also encoded a 3C protease site between the C-terminus of the TCR ectodomains and the GCN4 zippers to allow for cleavage of zippers. IE^k^ MHC was cloned into pAcGP67A with acidic/basic zippers as described for TCRs. IE^k^ α and 2B4 α chain also encoded ybbr-tag sequence for direct protein labelling. The IE^k^β construct was modified with an N-terminal extension containing either the 2A peptide via a Gly-Ser linker or CLIP peptide via a Gly-Ser linker containing a thrombin cleavage site. Proteins were transiently expressed in High Five insect cells (BTI-TN-5B1-4) and purified using His-tag/Nickel according to published protocols [Birnbaum, et al., 2014].

For fluorescent labeling of SNAP-tagged proteins, 10 μM protein was incubated with 20 μM benzylguanine functionalized dye (New England Biolabs) in HBS buffer (50 mM HEPES, 150 mM NaCl, 1 mM TCEP, pH 7.4) for 1 h at room temperature or overnight on ice. For PD-L1 and TCR 10 μM protein was incubated with 30 μM tetramethylrhodamine-5-maleimide in HBS buffer for 1 h at room temperature. Excess dyes were removed using Zeba Spin Desalting Columns (ThermoFisher, 89882).

### Preparation of SNAP-DNA tethers

Oligonucleotides were ordered from IDT with a 3’/5’ terminal amine and labeled with BG-GLA-NHS as previously described [Farlow, et al., 2013]. The adhesion strands used in this study consisted of a 3’ 20mer region (5’-ACTGACTGACTGACTGACTG-3’) with a 5’ 80mer poly-dT and the complementary sequence (5’- CAGTCAGTCAGTCAGTCAGT-3’) also with a 5’ 80mer poly-dT. Conjugation to benzylguanine was performed as described [Farlow, et al., 2013]. His_10_-tagged SNAP was labeled at a concentration of 5 μM with a 3-fold excess of BG-DNA in HBS (50 mM HEPES, 150 mM NaCl and 1 mM TCEP, pH 7.4).

### Electroformation of giant unilamellar vesicles

Lipids were mixed with a molar composition of 94.9% POPC, 5% DGS-NTA, 0.1% DOPE-390 in chloroform (Electron Microscopy Sciences, 12550) and dried under vacuum for 1 h to overnight. Electroformation was performed in 370 mM sucrose according to published protocols [Schmid, et al., 2015]. GUVs were stored at room temperature and imaged within one week.

### Preparation of supported lipid bilayers

Small unilamellar vesicles (SUVs) were prepared from a mixture of 95.5% POPC, 2% DGS-NGA-Ni, and 0.5% PEG5000-PE. The lipid mixture in chloroform was evaporated under argon and further dried under vacuum. The mixture was then rehydrated with phosphate buffered saline pH 7.4 and cycled between -80°C and 37°C 20 times, and then centrifuged for 45 min at 35,000 RCF. SUVs made by this method were stored at 4°C and used within two weeks of formation. Supported lipid bilayers were formed in freshly plasma cleaned custom PDMS chambers on RCA cleaned glass coverslips. 100 μL of SUV solution containing 0.5 to 1 mg/ml lipid was added to the coverslips and incubated for 30 min. Unadsorbed vesicles were removed and bilayers were blocked by washing three times with reaction buffer (50 mM HEPES, 150 mM NaCl, 1 mM TCEP, 1 mg/mL bovine serum albumin, pH 7.4), and incubating for 20 min.

### Optical setup for spinning disk, total internal reflection fluorescence, and scanning angle interference microscopy

Imaging was performed on one of two Nikon TI-E microscopes equipped with a Nikon 60x Plan Apo VC 1.20 NA water immersion objective, or a Nikon 100x Plan Apo 1.49 NA oil immersion objective, and four laser lines (405, 488, 561, 640 nm), either a Hamamatsu Flash 4.0 or Andor iXon EM-CCD camera, and μManager software [Edelstein, et al., 2010]. A polarizing filter was placed in the excitation laser path to polarize the light perpendicular to the plane of incidence. Angle of illumination was controlled with either a standard Nikon TIRF motorized positioner or a mirror moved by a motorized actuator (Newport, CMA-25CCCL). Scanning angle microscopy was performed and analyzed as previously described [Carbone, et al., 2016].

### Reconstitution of membrane interfaces

Proteins were diluted in reaction buffer (50 mM HEPES, 150 mM NaCl, 1 mM TCEP, 1 mg/mL bovine serum albumen, pH 7.4) and then mixed 2:1 with GUVs, or added to supported lipid bilayers. 1 h after mixing, supported lipid bilayers were washed three times with reaction buffer and GUVs were added. Rapamycin (Sigma, R0395) was added to FRB-FKBP reactions at a final concentration of 5 μM. GUVs were allowed to settle for 30-60 min prior to imaging.

### Image analysis

Images were analyzed using FIJI software [Schindelin, et al., 2012]. To calculate percent exclusion values, regions of interest (ROIs) were selected from protein of interest enriched and depleted zones of the GUV-SLB interface. The ratio of the mean pixel values for each GUV zone was calculated. This ratio was expressed as a percent corresponding to (1-(depleted/enriched))^*^100.

### Liposome Assay

Experiments were carried out as previously described [Hui et al, 2017]. Briefly, proteins were purified using baculovirus or bacterial expression system. LUVs and proteins of interest were premixed and incubated at room temperature for 1 h. 2 mM ATP was then injected and rapidly mixed to trigger Lck mediated phosphorylation of CD3*ζ* and PD-1. 20 minutes after ATP addition, apyrase was added (t = 0 min) and the reactions were allowed to continue at room temperature. Equal fractions of the reactions were removed and terminated with SDS sample buffer at the indicated time points. Anti-phosphotyrosine antibody (pY20, Santa Cruz Biotechnology #SC-508) was used to detect phosphorylation by western blotting.

